# Genomic and phenotypic trait variation of the opportunistic human pathogen *Aspergillus flavus* and its non-pathogenic close relatives

**DOI:** 10.1101/2022.07.12.499845

**Authors:** E. Anne Hatmaker, Manuel Rangel-Grimaldo, Huzefa A. Raja, Hadi Pourhadi, Sonja L. Knowles, Kevin Fuller, Emily M. Adams, Jorge D. Lightfoot, Rafael W. Bastos, Gustavo H. Goldman, Nicholas H. Oberlies, Antonis Rokas

**Affiliations:** Department of Biological Sciences, Vanderbilt University, Nashville, TN, USA; Evolutionary Studies Initiative, Vanderbilt University, Nashville, TN, USA; Department of Chemistry & Biochemistry, University of North Carolina at Greensboro, Greensboro, NC, USA; Department of Microbiology and Immunology, University of Oklahoma Health Science Center, Oklahoma City, OK, USA; Biosciences Center, Federal University of Rio Grande do Norte, Natal-RN, Brazil; Faculdade de Ciências Farmacêuticas de Ribeirão Preto, Universidade de São Paulo, Ribeirão Preto, Brazil

**Keywords:** *Aspergillus flavus*, genomes, phenotypic variation, secondary metabolites, fungal keratitis

## Abstract

Fungal diseases affect millions of humans annually, yet fungal pathogens remain understudied. The mold *Aspergillus flavus* is a causative agent of both aspergillosis and fungal keratitis infections, but species closely related to *A. flavus* are not considered clinically relevant. To study the evolution of *A. flavus* pathogenicity, we examined genomic and phenotypic traits of two strains of *A. flavus* and three closely related non-pathogenic species: *Aspergillus arachidicola* (two strains), *Aspergillus parasiticus* (two strains), and *Aspergillus nomiae* (one strain). We identified over 3,000 orthologous proteins unique to *A. flavus*, including seven biosynthetic gene clusters present in *A. flavus* strains and absent in the three non-pathogenic species. We chose to characterize secondary metabolite production for all seven strains under two clinically relevant conditions, temperature and salt concentration. Temperature impacted metabolite production in all species. Conversely, we found a lack of impact of salinity on secondary metabolite production. Strains of the same species produced different metabolites. Growth under stress conditions revealed additional heterogeneity within species. Using the invertebrate model of fungal disease *Galleria mellonella*, we found virulence of strains of the same species varied widely, and *A. flavus* strains were not more virulent than strains of the non-pathogenic species. In a murine model of fungal keratitis, we observed significantly lower disease severity and corneal thickness for *A. arachidicola* compared to other species at 48 hrs, but not at 72 hrs. Our work identifies key phenotypic, chemical, and genomic similarities and differences between the opportunistic human pathogen *A. flavus* and its non-pathogenic relatives.

## INTRODUCTION

Fungal infections affect millions of people worldwide annually but remain understudied and poorly understood (1). Severe fungal infections in humans, such as aspergillosis, commonly affect immunocompromised individuals (2). Invasive aspergillosis is an umbrella term describing a range of respiratory infections caused by inhalation of asexual spores of several *Aspergillus* species that grow into human tissue (3). Around 300,000 cases of invasive aspergillosis are identified each year, with a high mortality rate (2). Another form of aspergillosis, chronic pulmonary aspergillosis, impacts around three million people annually (2). Invasive aspergillosis and chronic pulmonary aspergillosis can also co-occur in immunocompetent patients with severe viral infections, such as SARS-CoV-2 (4) and influenza (5).

Fungal keratitis, or inflammation of the cornea due to a fungal infection, affects otherwise healthy individuals, although immunocompromised individuals have higher rates of infection than immunocompetent ones (2). Globally, at least 1,000,000 cases of fungal keratitis occur annually, including both yeast and filamentous fungal infections (2), over 15,000 of which are in the U.S, and ∼10% of cases result in the eye removal due to late diagnosis or poor therapeutic outcomes (6). Fungal keratitis can lead to blindness, and affected individuals are typically infected through small wounds on the eye’s surface (7). The disease primarily afflicts outdoor workers (e.g., farmers) and contact lens wearers (8, 9). Sand and vegetative material often cause the initial wounds through which the infection spreads (10). In western countries, including in the U.S., contact lens use is the primary risk factor (11). The Global Action Fund for Fungal Infections has designated fungal keratitis a public health priority (12). Childhood fungal keratitis is the most frequent cause of corneal blindness worldwide (13), which occurs in 40% of severe fungal keratitis cases (12).

*Aspergillus flavus*, an opportunistic human pathogen, is estimated to cause 10% of invasive aspergillosis infections worldwide, second only to *Aspergillus fumigatus*, and up to 80% of keratitis infections from *Aspergillus* species (14). Retrospective studies from Cuba and Pakistan identified more cases of chronic pulmonary aspergillosis caused by *A. flavus* than other *Aspergillus* species (15, 16). *A. flavus* is widespread in the environment, particularly in tropical regions (17), but we do not currently know which *A. flavus* traits impact virulence in humans. The taxonomic section *Flavi* includes *A. flavus* and closely related species such as *A. parasiticus*, *A. arachidicola*, and *A. nomiae* (18). Species in section *Flavi* share several morphological characteristics (19), including production of the carcinogen aflatoxin (18), but do not cause human disease at similar rates (20). For example, *A. flavus* is more commonly isolated in invasive aspergillosis and fungal keratitis cases than its close relative *A. parasiticus* (17). Interestingly, the two species share many characteristics. For example, *A. flavus* and *A. parasiticus* exhibit similar geographic ranges and both grow well at 37°C and 42°C (21); thermotolerance is considered key for *Aspergillus* pathogenicity in general (22, 23). Growth at 37°C has also been shown to change the metabolic profile of *Aspergillus* species (24). Secondary metabolites, which are small organic molecules with potent bioactivities, enable fungi to augment their environment with defensive compounds (25) and can impact virulence by modulating host biology (26, 27), as with gliotoxin in *A*. *fumigatus* (28). *A. flavus* and other section *Flavi* species have an even larger arsenal of secondary metabolites than *A. fumigatus* (29). Temperature can also impact secondary metabolite production. At 37°C, *A. flavus* produces diverse secondary metabolites (30). At lower temperatures, the most notable *A. flavus* secondary metabolite is the mycotoxin aflatoxin, a carcinogen (30), which is also produced by several other section *Flavi* species, including *A. parasiticus* (18). *A. flavus* is also predicted to produce scores of other secondary metabolites (31), which may play a role in human infection.

Current knowledge of *A. flavus* molecular mechanisms involved in human virulence rely heavily on extrapolation from studies of an *A. fumigatus* mouse model of both fungal keratitis (32) and respiratory infections (33) rather than direct studies of *A. flavus*, despite extensive genomic and phenotypic differences between the two species. Although in the same genus, *A. flavus* and *A. fumigatus* are not close relatives and are in distinct taxonomic sections of *Aspergillus*, sections *Flavi* and *Fumigati*, respectively (18); at the level of genome sequence divergence, the two species are as similar to each other as humans are to fish (34). Compared to *A. fumigatus*, section *Flavi* species have, on average, larger genomes, encode more genes, and are predicted to contain more biosynthetic gene clusters (BGCs) involved in the biosynthesis of secondary metabolites (35). Differences between *A. flavus* and *A. fumigatus* include resistance to antifungal drugs, as azole resistance differs between the two species, with triazole considered resistance rare in *A. flavus* despite comparable exposure to environmental fungicides as *A. fumigatus* (36). Additional studies investigating genetic determinants of virulence specific to *A. flavus*, which enable the species to infect humans at a higher rate than closely related species, as well as the drug resistance profiles of diverse strains and species, are needed to better understand the pathogenicity of *A. flavus*.

To begin addressing the evolution of pathogenicity in section *Flavi*, we examined the genomes, secondary metabolite profiles, as well as other infection-relevant phenotypic traits (e.g., virulence) of two *A. flavus* strains and strains from three closely related species considered non-pathogenic: *A. arachidicola* (two strains), *A. parasiticus* (two strains), and *A. nomiae* (one strain). We sequenced seven strains from the four species and identified proteins unique to *A. flavus* strains, including BGCs absent in the non-pathogenic species. Temperature impacted secondary metabolite production in all four species, but secondary metabolites unique to *A. flavus* were not identified under these conditions. Antifungal drug resistance did not differ appreciably between strains or species, but strains of the same species exhibited variation in growth under certain stress conditions. Evaluation of virulence using an invertebrate model of disease revealed additional variation between strains of the same species, and *A. flavus* strains were not the most virulent in this model. Virulence in a murine model of fungal keratitis was tested with one strain each from *A. flavus*, *A. parasiticus*, and *A. arachidicola*, along with a reference strain of *A. fumigatus*, revealing significantly lower disease severity and corneal thickness in mice infected with *A. arachidicola* compared to the other species. The combination of genomic and phenotypic comparisons revealed similarities and differences between *A. flavus* and three non-pathogenic close relatives within *Aspergillus* section *Flavi*. This study provides key data on the genomic, chemical, and phenotypic diversity of closely related pathogenic and non-pathogenic species in *Aspergillus* section *Flavi* that further our knowledge of how some of these fungi can opportunistically infect humans.

## MATERIALS AND METHODS

### Genome sequencing and comparisons

#### 1. Strains and growth conditions

Seven strains from four *Aspergillus* species were obtained from the USDA Agricultural Research Service Culture Collection (*A. flavus* NRRL 501, *Aspergillus nomiae* NRRL 6108, and *Aspergillus parasiticus* NRRL 2999) or from the laboratory of Dr. Ignazio Carbone at North Carolina State University (*Aspergillus flavus* NRRL 1957, *Aspergillus parasiticus* NRRL 502, and *Aspergillus arachidicola* IC26645 and IC26646). All strains were maintained on Potato Dextrose Agar (PDA; Difco) at room temperature (approximately 23°C).

#### 2. Genomic DNA extraction and sequencing

For genomic DNA extraction to facilitate genome sequencing, the strains were first grown on PDA (Difco). Each strain was subsequently transferred to a 100 mm Petri dish with an overlay of sterile autoclaved Hybond nylon membranes (84 mm) obtained from Amersham (GE Healthcare). The strains were then allowed to grow for 2-3 weeks on PDA media overlaid with this membrane. After suitable growth was achieved, a sterile scalpel was used to harvest the fungal mycelia by scraping the surface of the nylon membrane without dislodging any agar from the Petri plate. Using a sterile mortar and pestle, the mycelia were then ground to a fine powder with liquid nitrogen, transferred to a bashing bead tube with 750 µl of DNA lysis buffer (Zymo Quick-DNA fungal/bacterial miniprep kit). The powder in the bashing bead tube was further disrupted and homogenized in a Qiagen TissueLyser LT bead mill for 5 min. Genomic DNA was obtained using the standard protocol in the Zymo Quick-DNA fungal/bacterial miniprep kit booklet.

Paired-end sequencing (2 x 150 bp) of the gDNA was performed at the Vanderbilt Technologies for Advanced Genomics (VANTAGE) facility using the NovaSeq 6000 platform (Illumina, Inc.) following manufacturer’s protocols. Libraries were prepared using the Illumina TruSeq DNA PCR-free kit.

#### 3. Genome assembly, annotation, and assessment

Adaptors were removed from the reads and filtered for quality using Trimmomatic v0.39 (37). Trimmed reads were assembled into draft genomes using SPAdes v3.12.0 (38) with the “--careful” flag. Scaffolds under 500 bp were removed from further analysis. Draft genomes were annotated using Liftoff v1.2.0 (39), a sequence-similarity method which requires a reference annotation. *A. flavus* strains and *A. nomiae* NRRL 6108 were annotated using strain NRRL 3357 (40); for *A. arachidicola* strains we used CBS 117612 (29); *A. parasiticus* strains were annotated from CBS 117618 (29). The *A. parasiticus* NRRL 2999 draft genome was compared to that of *A. parasiticus* SU-1 to confirm isolate identity using fastANI (41), which compares average nucleotide identity between genomes. Genome and annotation completeness was evaluated using BUSCO v4.0.4 (42), compared against the “eurotiales” database of universal single-copy orthologs.

#### 4. Phylogenetics

In addition to the 7 newly sequenced genomes, the genomes and predicted proteomes of 23 other *Aspergillus* strains were downloaded from NCBI (Table 1). In total, we used 29 strains from 18 species of *Flavi* to build a species tree phylogeny of *Aspergillus* section *Flavi* with *Aspergillus niger* CBS 513.88 (section *Nigri*) serving as an outgroup. Single-copy orthologous proteins in all 30 strains were identified using OrthoFinder v2.5.4 (43). For each ortholog, all 30 copies were aligned using Muscle v3.8.1551 (44) and the amino acid alignments trimmed using trimAl v1.2 (45) using the “gappyout” option. Alignments shorter than 200 amino acids were removed from analysis and the remaining alignments were concatenated into a single data matrix using the script catfasta2phyml.pl (https://github.com/nylander/catfasta2phyml). A maximum likelihood phylogeny was reconstructed using IQ-tree v1.6.1 (46) with the “JTT+F+I+G4” model and 1,000 replicates for bootstrapping. The resulting consensus tree was viewed using iTOL (47).

**Table 1.**
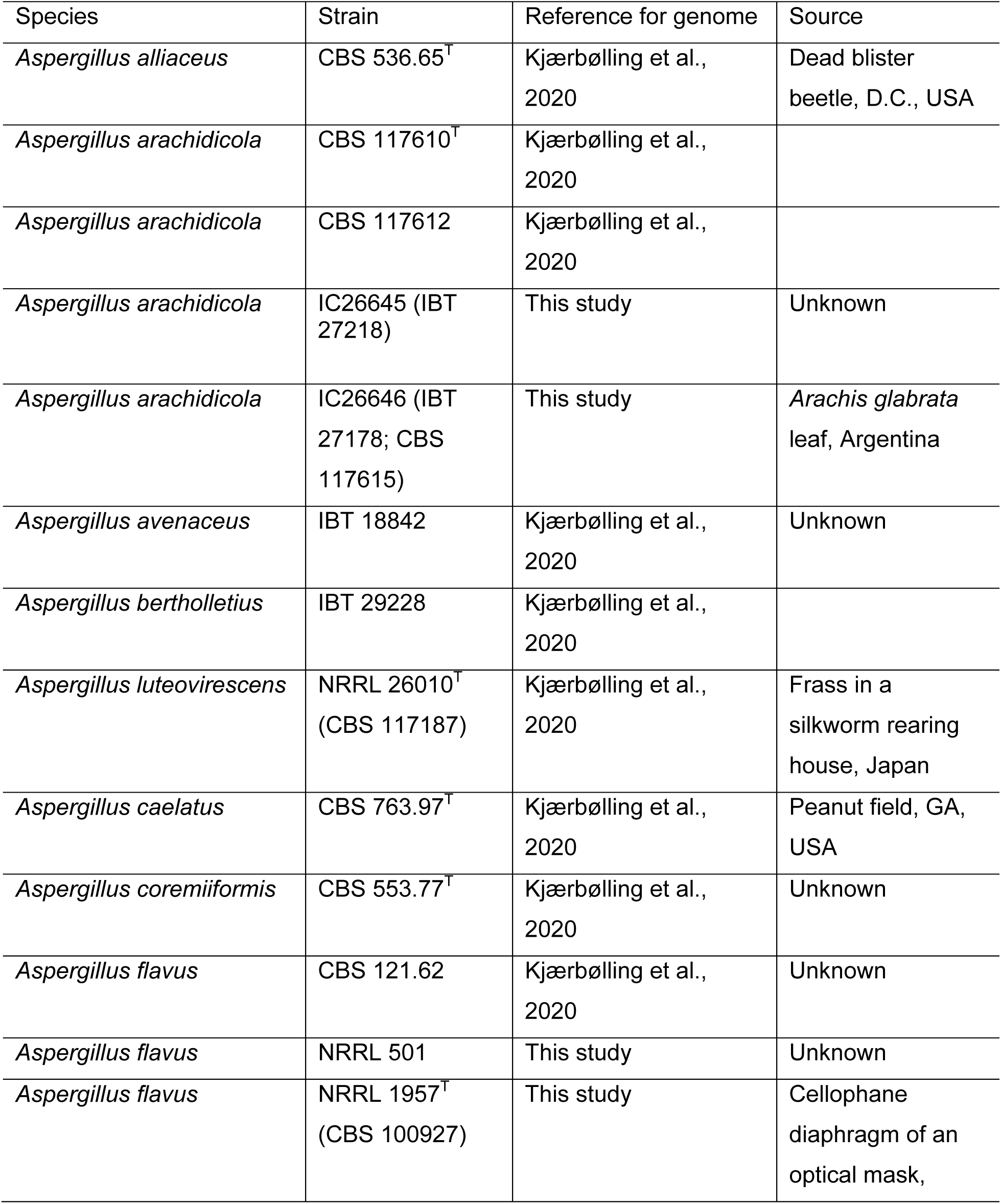
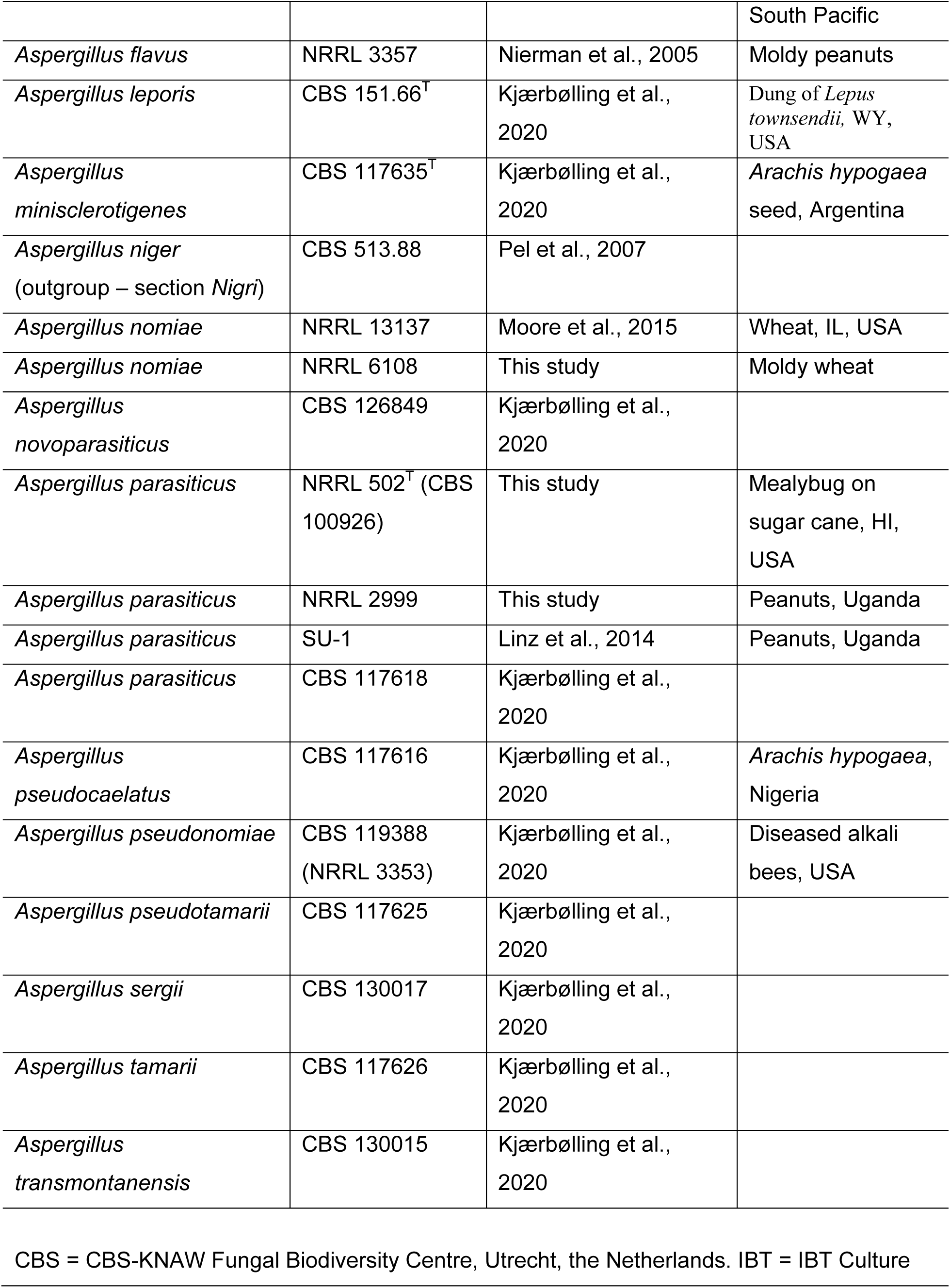
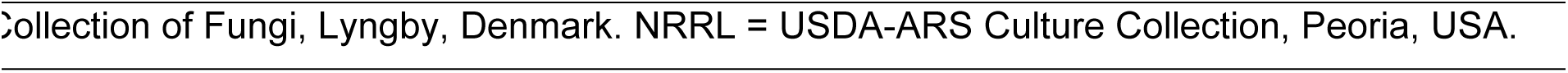
Genomes of 27 Aspergillus strains from 20 species including 7 newly sequenced genomes.

#### 5. Identification of orthologous and unique protein families and biosynthetic gene clusters

Using the seven newly sequenced strains (*A. flavus* NRRL 501 and NRRL 1957, *A. parasiticus* NRRL 502 and NRRL 2999, *A. nomiae* NRRL 6108, and *A. arachidicola* IC26645 and IC26646), we compared each predicted proteome in a pairwise manner and in combination to identify unique and orthologous proteins for all combinations using OrthoVenn2 (48). Within the unique or orthologous protein groups, enriched gene ontology (GO) categories were identified from UniProt annotations (49) and probability testing was based on hypergeometric distribution (48). The R package “UpSetR” was used to visualize the number of shared protein families in each grouping as an upset plot. Biosynthetic gene clusters (BGCs) were predicted using fungiSMASH v6.0 (50). BGCs from different strains were compared using BiGSCAPE-CORASON (51) with cutoff set to 0.70, and BGCs shared between all strains and strains of the same species were identified. Synteny plots of BGCs of interest were visualized using Clinker (52).

### Secondary metabolite isolation and structural elucidation

The profile of secondary metabolites for the seven *Aspergillus* section *Flavi* strains were investigated using well established procedures (24). Each strain was grown at both 23°C (i.e., room temperature) and 37°C on oatmeal (Old fashioned breakfast Quaker oats). Since the lacrimal fluid is composed of physiologic saline, the oatmeal cultures were prepared both with and without 6 mg/ml saline to assess if saline impacted the profile of secondary metabolites.

Solid-state fermentations (n = 6) were carried out in broad mouth 250 ml Erlenmeyer flasks. To start oatmeal cultures, an agar plug from the leading edge of a PDA Petri dish culture was transferred to a sterile tube with 10 ml of liquid YESD (YESD; 20 g soy peptone, 20 g dextrose, 5 g yeast extract, 1 L distilled H_2_O) and grown for seven days on an orbital shaker (100 rpm) at room temperature (∼23°C) and used to inoculate the oatmeal solid fermentation media. Oatmeal cereal media (Old fashioned breakfast Quaker oats) were prepared by adding 10 g oatmeal to a 250 mL Erlenmeyer flask with either 15-17 mL of DI-H_2_O or 15-17 mL of a 6 mg/ml saline solution (i.e., 6 g of Instant Ocean in 1000 ml of DI-H_2_O), then autoclaved at 121°C for 30 min. For each of the seven strains, six fermentation flasks were incubated at room temperature with saline and six without saline; similarly, six flasks were incubated at 37°C with saline and six without saline. Therefore, for each strain, 24 Erlenmeyer flasks were grown in total. Prior to analysis of the secondary metabolite profile, the cultures were incubated statically at room temperature for 14 days or 37°C for 7 days.

#### 1. Extraction of secondary metabolites

For each condition, each of the six culture flasks was extracted individually and treated as a biological replicate. Each individual flask was extracted by adding 60 ml of CHCl_3_-MeOH (1:1), chopping with a spatula, and shaking overnight (**≈** 16 h) at 100 rpm at room temperature. The cultures were then filtered *in vacuo*, and 90 ml of CHCl_3_ and 150 ml of DI-H_2_O were added to each of the filtrates. The mixture was then transferred to a separatory funnel and shaken vigorously. The organic layer (i.e., bottom layer) was drawn off and evaporated to dryness *in vacuo*. The dried organic layer was reconstituted in a 100 ml mixture of CH_3_CN-MeOH (1:1) and 100 ml of hexanes, transferred to a separatory funnel, and shaken vigorously. The defatted organic layers (i.e., CH_3_CN-MeOH layer) were evaporated to dryness.

All the defatted organic layers were analyzed individually by UPLC-HRMS utilizing a Thermo LTQ Orbitrap XL mass spectrometer equipped with an electrospray ionization source. A Waters Acquity UPLC was utilized using a BEH C_18_ column (1.7 μm; 50 mm × 2.1 mm) set to a temperature of 40°C and a flow rate of 0.3 mL/min. The mobile phase consisted of a linear gradient of CH_3_CN−H_2_O (both acidified with 0.1% formic acid), starting at 15% CH_3_CN and increasing linearly to 100% CH_3_CN over 8 min, with a 1.5 min hold before returning to the starting conditions.

#### 2. Metabolomic analysis

Principal component analysis (PCA) and hierarchical clustering were performed on the UPLC-HRMS data. Untargeted UPLC-HRMS datasets for each sample were individually aligned, filtered, and analyzed using Mzmine 2.53 software (https://sourceforge.net/projects/mzmine/). Peak list filtering and retention time alignment algorithms were used to refine peak detection, and the join algorithm integrated all sample profiles into a data matrix using the following parameters: Mass detection: MS1 positive mode. ADAP: Group intensity threshold: 20000, Min highest intensity: 60000, *m/z* tolerance: 0.003. Chromatogram deconvolution: Wavelets (ADAP) algorithm. Join aligner and gap filling: retention time tolerance: 0.05 min, m/z tolerance: 0.0015 *m/z*.

The resulting data matrix was exported to Excel (Microsoft) for analysis as a set of *m/z*– RT (retention time) pairs with individual peak areas. Samples that did not possess detectable quantities of a given marker ion were assigned a peak area of zero to maintain the same number of variables for all sample sets. Ions that did not elute between 1 and 10 min and/or had an *m/z* ratio < 200 or > 900 Da were removed from analysis. Relative standard deviation was used to understand the quantity of variance between the injections, which may differ slightly based on instrument variance. A cutoff of 1.0 was used at any given *m/z*–RT pair across the biological replicate injections, and if the variance was greater than the cutoff, it was assigned a peak area of zero. PCA analysis and hierarchical clustering were created with Python. The PCA scores plots were generated using the averaged data of the six individual biological replicates.

#### 3. Isolation and identification of secondary metabolites

After comparison of the UPLC-HRMS data, the defatted organic layers for each condition were combined due to the similarity of their chemical profiles, so as to generate a larger pool of material for isolation studies, which were carried out using well established natural products chemistry procedures (53, 54). The fractions were dissolved in CHCl_3_-MeOH, absorbed onto Celite 545 (Acros Organics), and fractioned by normal phase flash chromatography using a gradient of hexane-CHCl_3_-MeOH. The isolation of the compounds was carried out using preparative HPLC.

The isolated fungal metabolites were identified by direct comparison of the spectroscopic and spectrometric properties with those previously reported, and where possible, structures were validated by comparisons with authentic reference standards (55). Additionally, mass defect filtering (56) was used to identify structurally-related analogues of the isolated compounds.

#### 4. General experimental procedures

The NMR data were collected using a JEOL ECS-500 spectrometer operating at 500 MHz for ^1^H and 125 MHz for ^13^C or an Agilent 700 MHz spectrometer, equipped with a cryoprobe, operating at 700 MHz for ^1^H and 175 MHz for ^13^C. The HPLC separations were performed on a Varian Prostar HPLC system equipped with a Prostar 210 pump and a Prostar 335 photodiode array detector (PDA), with the collection and analysis of data using Galaxy Chromatography Workstation software. The columns used for separations were either a Synergi C_18_ preparative (4 μm; 21.2 × 250 mm) column at a flow rate of 21.2 mL/min, Luna PFP(2) preparative (5 μm; 21.2 × 250 mm) column at a flow rate of 17 mL/min or an Atlantis T3 C_18_ preparative (5 μm; 19 × 250 mm) column at a flow rate of 17 mL/min. Flash chromatography was performed on a Teledyne ISCO Combiflash Rf 200 as monitored by ELSD and PDA detectors.

### Growth assays

#### 1. Antifungal drug susceptibility testing

Antifungal susceptibility testing for both voriconazole (Sigma-Aldrich), and amphotericin B (Sigma-Aldrich) was performed by determining the minimal inhibitory concentration (MIC) according to the protocol established by the Clinical and Laboratory Standards Institute (CLSI, 2017).

#### 2. Stress response

Radial growth was used to compare how the different strains respond to cell wall stress, iron starvation, and hypoxia. Strains were grown in solid minimal medium (MM), as described previously (57), inoculated with 1X10^5^ spores of each strain, and incubated for 5 days at 37°C before colony diameter was measured. To induce cell wall stress, 30 and 50 μg/mL of Congo red (cell wall perturbing) was added to medium. For iron starvation, iron-poor MM was devoid of all iron and supplemented with 128 μg/mL of Gallium Nitrate (Sigma-Aldrich). Gallium is chemically similar to iron, and for this reason it is taken up by the cell replacing iron (58). Radial growth for the aforementioned stresses was expressed as ratios, dividing colony radial diameter (cm) of growth in the stress condition by colony radial diameter in the control (no stress) condition. For hypoxia analysis, the plates were incubated on 5% CO_2_ and 1% O_2_ at 37°C for 5 days.

Oxidative stress was measured using the protocol described by Canóvas et al., (59). Briefly, the experiments were performed in 96-wells plates containing 100 μL of MM (with 1% agar) supplemented or not with 3mM of H_2_O_2_ (Merck S.A) or 0.15mM of menadione (Sigma-Aldrich). Each well was inoculated with 1X10^5^ spores, and the growth was measured over time by quantifying the absorbance at 595 nm in a plate reader (Synergy HT) at 37°C. Data were recorded and analyzed with Gen5TM Data Analysis Software v2.0 and exported to Microsoft Excel for further analysis and generation of the graphs. Lag times were calculated using Gen5TM Data Analysis Software v2.0 as the time interval between the line of maximum slope of the propagation phase and the absorbance baseline at time zero.

#### 3. Statistical evaluation of growth assays

A one-way ANOVA was calculated for the iron starvation, hypoxia, and oxidative stress datasets and a two-way ANOVA was calculated for the dataset of cell wall stress with two Congo red concentrations. Graphs were visualized and statistics calculated using GraphPad Prism v9.3.1.

### Assessment of virulence using the invertebrate model of fungal disease Galleria mellonella

#### 1. Preparation of larvae and inoculum

*Galleria mellonella* larvae were used to investigate the virulence of all seven strains. The larvae used for the infection were in the last larval stage of development (sixth week). All selected larvae weighed ∼300 mg and were restricted to food for 24 h before the experiment. Fresh asexual spores (conidia) of each strain were counted using a hemocytometer.

#### 2. Inoculation and observation

Five uL of each inoculum were injected using a Hamilton syringe (7000.5 KH) through the last left ear (n = 10/group), resulting in 1 × 10^6^, 1 × 10^4^, and 1 × 10^3^ conidia/larvae. The control group was inoculated with phosphate buffered saline (PBS). After infection, the larvae were kept with food restrictions, at 37°C in Petri dishes in the dark and scored daily for fifteen days. The larvae were considered dead due to lack of movement in response to touch. The viability of the inoculum administered was determined by serial dilution of the conidia in YAG medium and incubating the plates at 37°C for 72 h. The experiment was repeated twice.

We separated and assembled the groups with the larvae (n = 10) in Petri dishes. The groups are composed of larvae that are approximately 300 mg in weight and 2 cm long. Moth sex was not accounted for due to the impossibility of visually determining sex at the sixth week of larval development.

#### 3. Statistical analysis of infection rates

Larval survival was plotted on Kaplan-Meier curves using the “survival” and “survminer” R packages. A Mantel-Cox log rank test was used to evaluate statistical significance between the survival curves of larvae infected with different fungal strains.

### Assessment of virulence using a murine model of keratitis

#### 1. Preparation of the fungal inoculum

On the day of corneal inoculation, asexual spores (conidia) of *A. arachidicola* IC26646, *A. flavus* NRRL 1957*, A. fumigatus* Af293, and *A. parasiticus* NRRL 2999 were incubated in 25 mL YPD broth (yeast extract, peptone, dextrose) at a density of 5 X 10^6^ conidia/ml at 35°C, 200 RPM. Once conidia were swollen and clumping, but not polarized (approximately 4h for all strains), cultures were collected by centrifugation, washed twice with PBS, resuspended in PBS to an optical density of 0.8 (360 nm), and stored at room temperature until the corneal inoculation (approximately 1 hr).

#### 2. Corneal infections

On the day preceding corneal inoculation, 6–8-week-old male C57BL/6J mice (Jackson Laboratories) were immunosuppressed with a 100 mg/kg methylprednisolone by intraperitoneal (i.p.) injection. The following day, animals were anesthetized with 100 mg/kg ketamine and 6.6 mg/kg xylazine, i.p., and the corneal epithelium was ulcerated over the pupil of the right eye to a diameter of approximately 1 mm using an Algerbrush II. Five µL of the fungal inocula (described above) were pipetted over the ulcerated corneas and remained in place for 20 minutes before being removed with a Kim wipe. Five animals were included in each infection group, where the contralateral eye of all animals remained uninfected in accordance with the Association for Research in Vision and Ophthalmology (ARVO) guidelines for the use of animals in vision research.

Animals were further injected subcutaneously with 1 mg/kg Buprenorphine SR for analgesia.

#### 3. Slit-lamp microscopy and disease scoring

Each day post-inoculation (p.i.), animals were anesthetized with isoflurane and imaged by slit-lamp using a Micron IV Biomicroscope (Phoenix Technology Group, CA, USA). Images were de-identified and assigned an overall disease score (range 0-4) by two blinded reviewers based on the area of opacification, density of opacification, and surface irregularity. The average disease scores for each cornea were compared by one-way ANOVA using GraphPad Prism v9.3.1.

#### 4. Optical coherence imaging and corneal thickness measurement

Corneas were also imaged using the Bioptigen spectral domain-optical coherence tomography (SD-OCT) system (Leica Microsystems, Deerfield, IL, USA). Mice were anesthetized with isoflurane and a 4 x 4 mm image was scanned with a 12 mm telecentric lens. Reference arm calibration was completed by the manufacturer and set to 885. Images were analyzed using the InVivoVue Diver software (Bioptigen). Briefly, corneal scans were digitally overlaid with a 11 x 11 spiderplot and the distance between the epithelium and endothelium was measured at 11 distinct points near the central cornea. The average of the 11 measures was taken as the corneal thickness (mm) and compared across groups using a one-way ANOVA in GraphPad Prism v9.3.1.

#### 5. Fungal burden assessment

At 72 h p.i., corneas were resected and homogenized by incubation in 1mL of buffer containing 2 mg/mL collagenase 1 (Sigma) for 1 h at 37°C. Dilutions of the homogenate were plated onto inhibitory mold agar, incubated overnight at 35°C, and colonies were enumerated. Colony counts were compared between groups using a one-way ANOVA in GraphPad Prism v9.3.1.

## RESULTS

### Draft genomes for seven strains of A. flavus and non-pathogenic close relatives

We sequenced and assembled seven genomes from four species with high coverage (81x – 230x). The *A. flavus* NRRL 1957 draft genome consisted of the fewest scaffolds (143), whereas the draft genome of *A. arachidicola* IC26646 had the most (1, 440). *A. nomiae* NRRL 6108 had the smallest genome, at 36.8 Mbp, and *A. parasiticus* NRRL 502 had the largest, at 41.4 Mbp; *A. flavus* strains had smaller genomes than strains of *A. parasiticus* or *A. arachidicola* (Table 2). All genomes had over 93% of the expected universal single-copy orthologs (Table 2).

**Table 2.** Genome sequencing, assembly, and annotation for seven strains of four Aspergillus species revealed expected genome size and good coverage depth.

| Species | Strain | Total paired reads | Trimmed paired reads | No. scaffolds | N50 | Coverage | Size (Mbp) | No. predicted proteins |
| --- | --- | --- | --- | --- | --- | --- | --- | --- |
| *Aspergillus arachidicola* | IC26645 | 30,016,585 | 28,186,702 | 805 | 1,139,563 | 102x | 41.2 | 13,710 |
| *Aspergillus arachidicola* | IC26646 | 32,284,138 | 29,911,439 | 1,440 | 1,298,300 | 109x | 40.9 | 13,731 |
| *Aspergillus flavus* | NRRL 1957 | 44,750,116 | 40,985,045 | 143 | 1,892,429 | 163x | 37.6 | 14,216 |
| *Aspergillus flavus* | NRRL 501 | 30,987,721 | 28,845,216 | 563 | 1,420,800 | 111x | 38.8 | 13,997 |
| *Aspergillus nomiae* | NRRL 6108 | 25,466,314 | 23,606,091 | 252 | 1,660,546 | 96x | 36.9 | 11,820 |
| *Aspergillus parasiticus* | NRRL 502 | 24,439,029 | 22,469,327 | 702 | 95,7843 | 81x | 41.5 | 13,470 |
| *Aspergillus parasiticus* | NRRL 2999 | 63,194,489 | 62,404,154 | 516 | 1,838,014 | 230x | 40.6 | 13,590 |

**Table 3.**
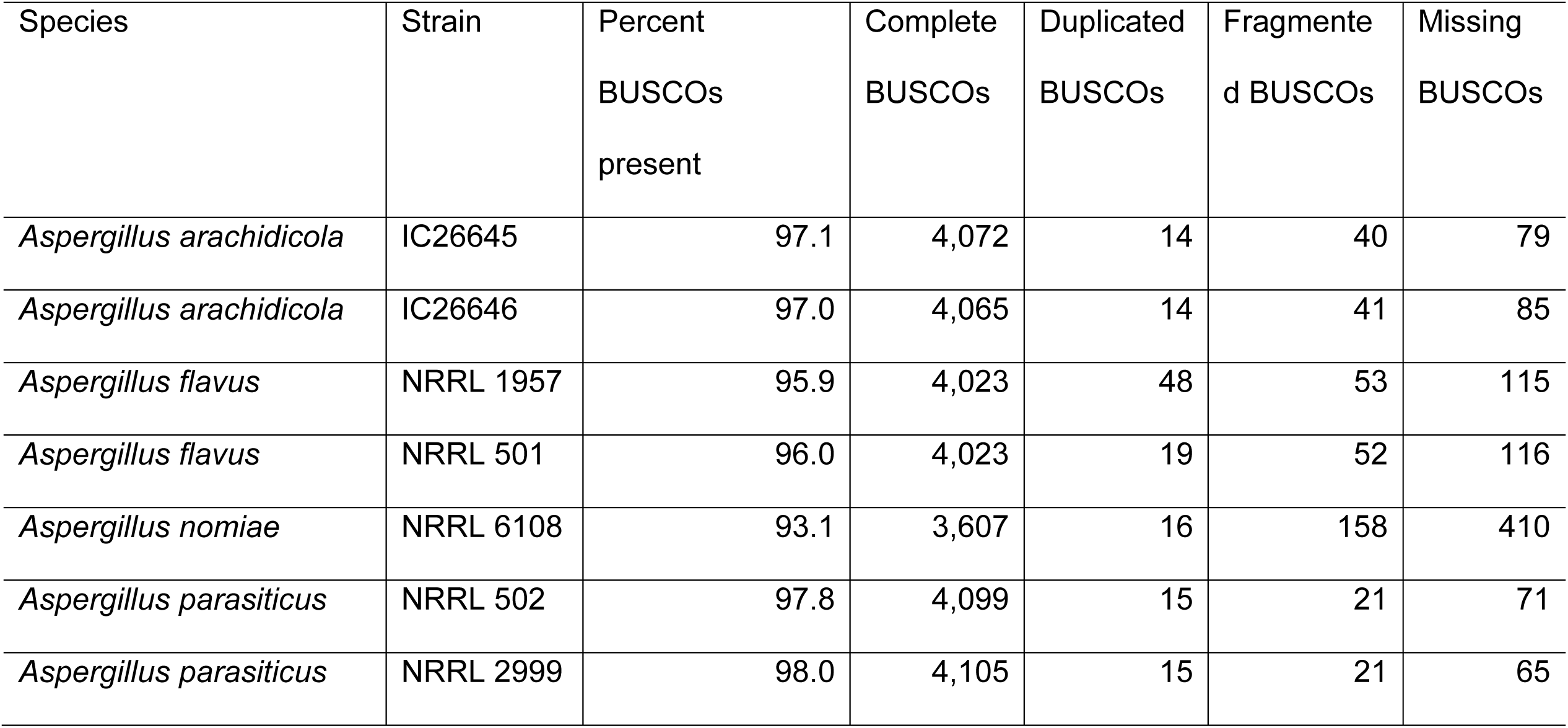
Genome completeness assessment of draft *Aspergillus* genomes sequenced for this study.

| Species | Strain | Percent<br>BUSCOs<br>present | Complete<br>BUSCOs | Duplicated<br>BUSCOs | Fragmente<br>d BUSCOs | Missing<br>BUSCOs |
| --- | --- | --- | --- | --- | --- | --- |
| *Aspergillus arachidicola* | IC26645 | 97.1 | 4,072 | 14 | 40 | 79 |
| *Aspergillus arachidicola* | IC26646 | 97.0 | 4,065 | 14 | 41 | 85 |
| *Aspergillus flavus* | NRRL 1957 | 95.9 | 4,023 | 48 | 53 | 115 |
| *Aspergillus flavus* | NRRL 501 | 96.0 | 4,023 | 19 | 52 | 116 |
| *Aspergillus nomiae* | NRRL 6108 | 93.1 | 3,607 | 16 | 158 | 410 |
| *Aspergillus parasiticus* | NRRL 502 | 97.8 | 4,099 | 15 | 21 | 71 |
| *Aspergillus parasiticus* | NRRL 2999 | 98.0 | 4,105 | 15 | 21 | 65 |

**Table 4.**
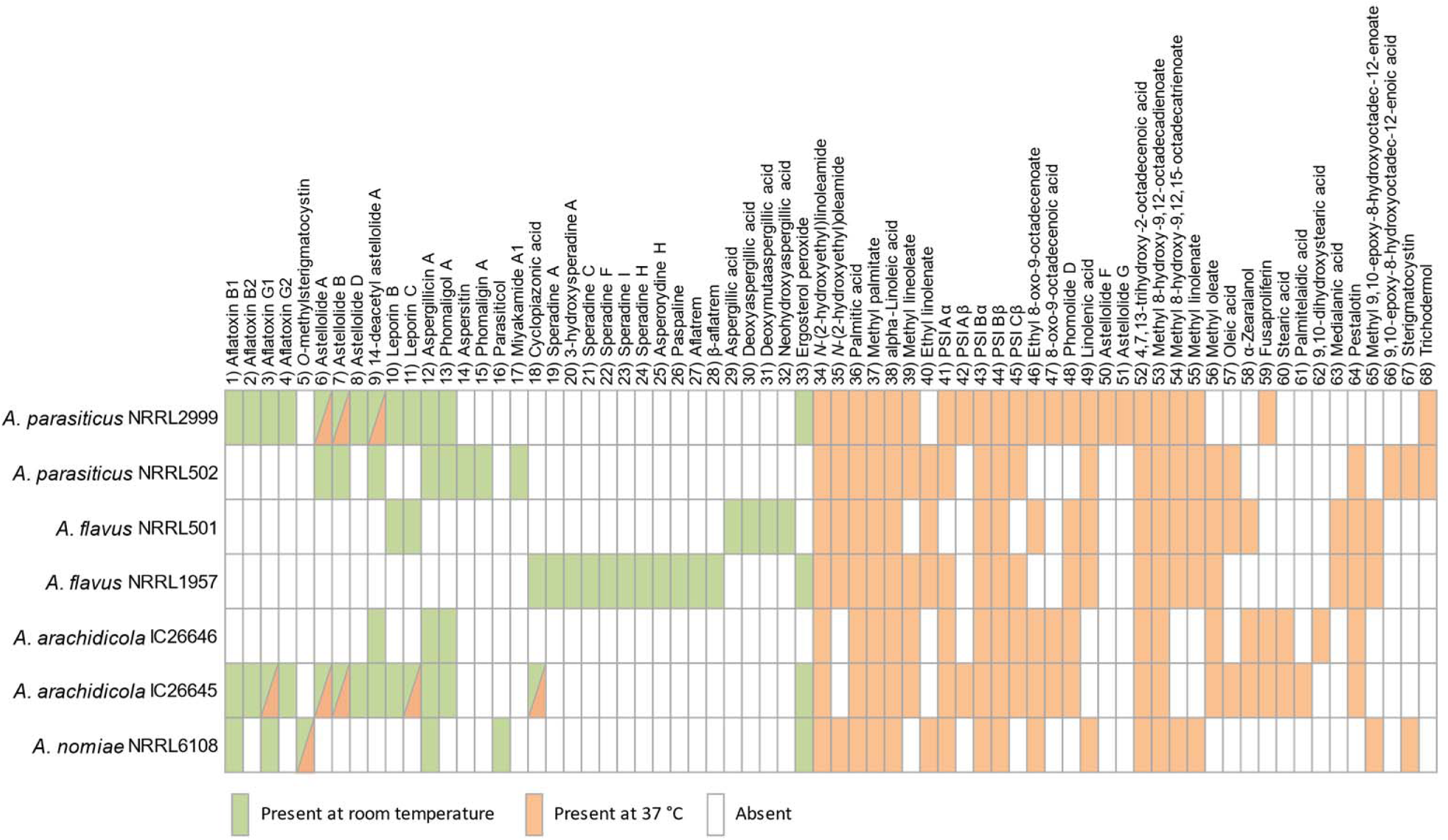
Variation in the presence or absence of secondary metabolites in seven strains from four Aspergillus species (green boxes represent metabolites produced at room temperature and orange boxes metabolites produced at 37°C)

### Placement of newly sequenced genomes consistent with Aspergillus section Flavi phylogeny

Our maximum likelihood species tree was built from 2,422 orthologs shared among 20 *Aspergillus* species (Fig. 1). Species identification was confirmed for all the newly sequenced strains (NRRL 501, 502, 1957, 2999, 6108 and IC26645 and IC26646) with high support (BS = 100) with each strain placed among the representative strains of the expected species. We note that previous whole genome sequencing of a strain labeled “NRRL 2999” (GCA_012897115.1) was recently determined to be *A. flavus*, rather than *A. parasiticus* (60), and was acknowledged as a clonal derivative of *A. flavus* NRRL 3357 (61). Our NRRL 2999 strain and *A. parasiticus* SU-1 share 99.98% average nucleotide identify, confirming that our strain, obtained from the NRRL culture collection, is *A. parasiticus*.

**Figure 1.**
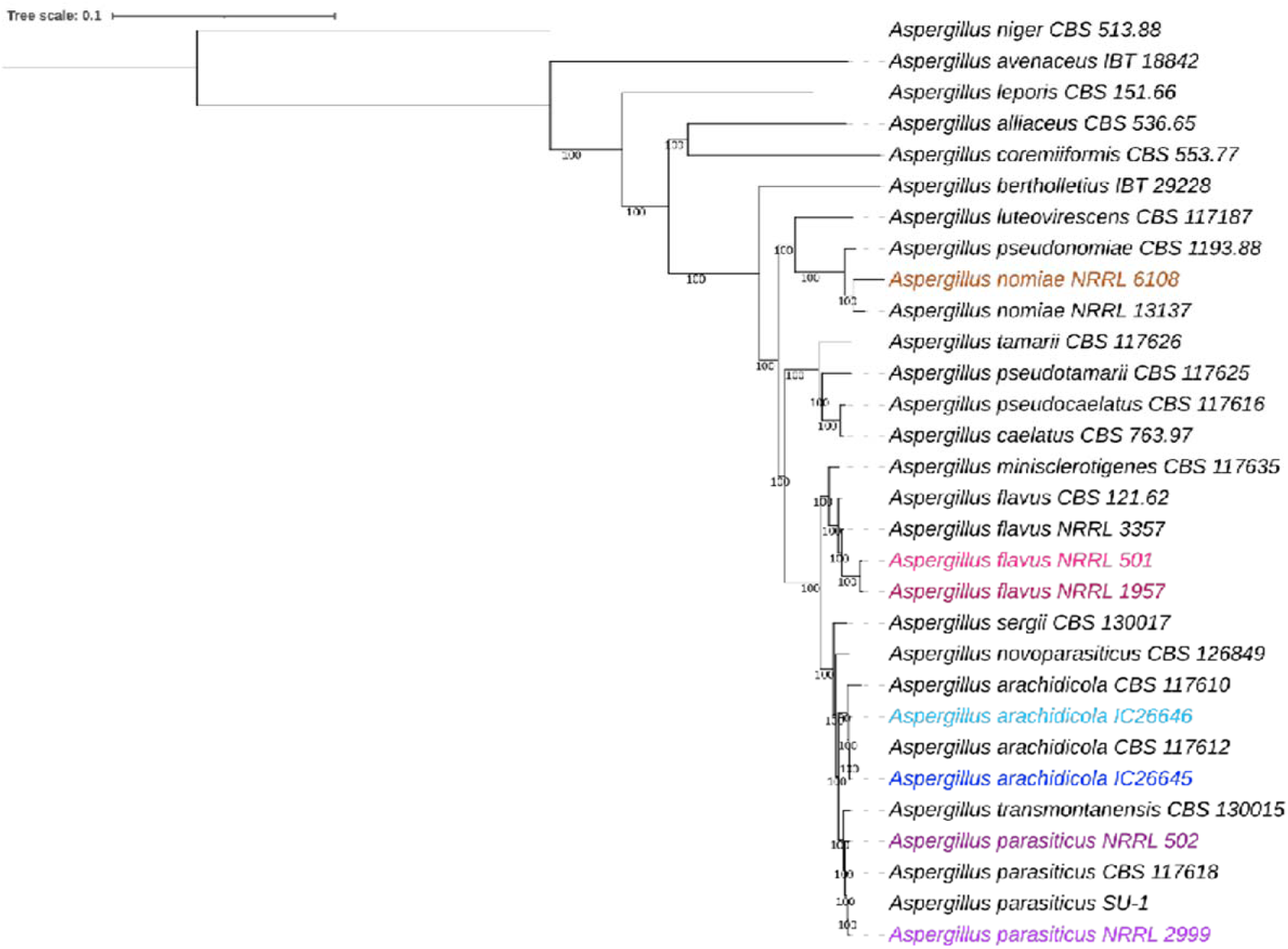
The taxonomic identity of newly sequenced strains is consistent with the *Aspergillus* section *Flavi* phylogeny. A maximum likelihood phylogeny was constructed using 2,422 single-copy orthologs from 30 strains of 20 *Aspergillus* species, using *A. niger* as the outgroup. Additional information about strains used can be found in Table 1. Numbers near branches are bootstrap values calculated from 1,000 replicates. Strains sequenced as part of this study are highlighted, with *A. nomiae* NRRL 6108 in orange, *A. flavus* in pink (NRRL 501 = light pink, NRRL 1957 = dark pink), *A. arachidicola* in blue (IC26645 = dark blue, IC26646 = light blue), and *A. parasiticus* in purple (NRRL 502 = dark purple, NRRL 2999 = light purple).

### More than 3,000 protein families are unique to the pathogen A. flavus

We identified 8,717 protein families shared by all seven newly sequenced strains (Fig. 2). A further 3,054 protein families were present in both *A. flavus* strains but absent from the other five strains. This group of protein families was enriched for the gene ontology (GO) terms GO:0016705 “oxidoreductase activity” (15 protein families, p < 0.0001, hypergeometric distribution) and GO: 0055085 “transmembrane transport” (30 protein families, p < 0.0001). *A. arachidicola* strains shared 3,097 unique protein families enriched for GO term GO:0016114 “terpenoid biosynthetic process” (10 protein families, p < 0.00001) and *A. parasiticus* strains shared 1,230 unique protein families without any GO term enrichment. No GO terms were enriched within the 1,322 protein families unique to *A. nomiae*.

**Figure 2.**
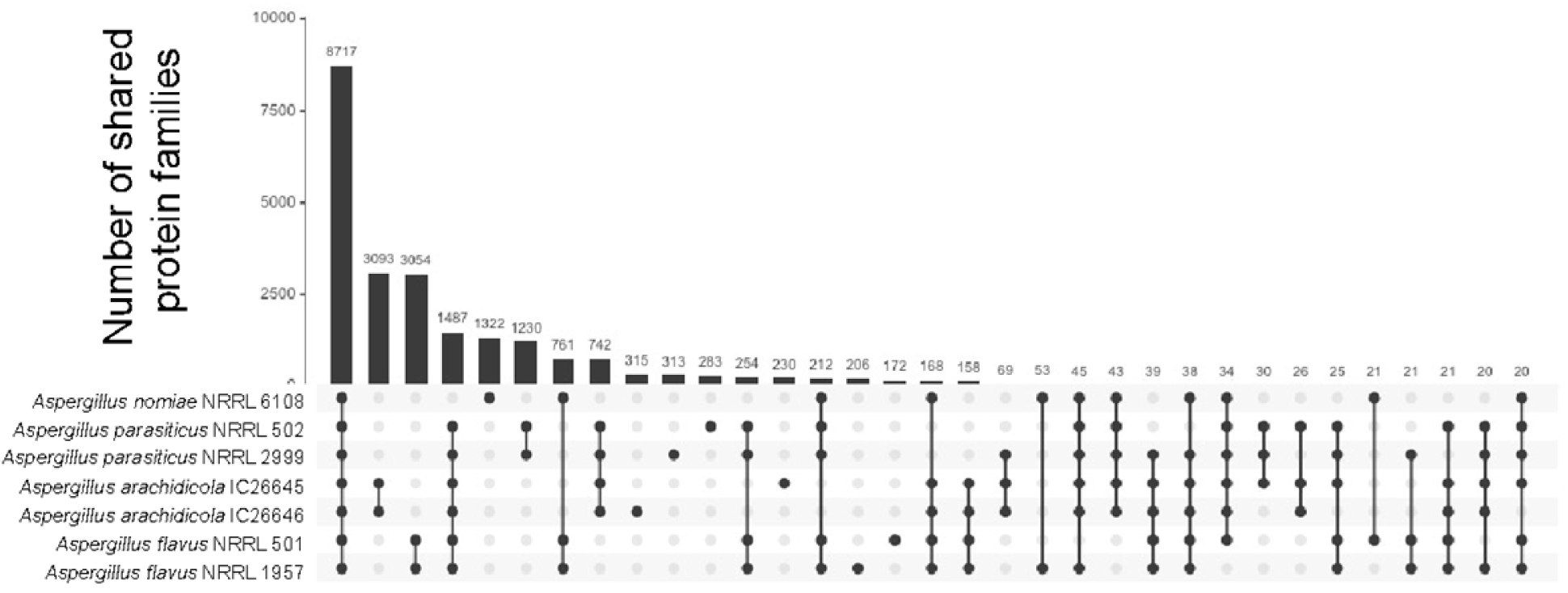
Strains of the same species do not differ substantially in predicted protein families. Upset plot showing number of shared protein families for orthogroups with at least 20 proteins. Linked black circles under bar plot indicate strains sharing the orthologous protein families. Numbers above bars indicate exact number of shared families.

### Many predicted biosynthetic gene clusters and secondary metabolites were shared between species

Using the fungal version of antiSMASH, we predicted biosynthetic gene clusters (BGCs) from the seven strains. *A. nomiae* NRRL 6108 had the fewest predicted BGCs at 49, whereas *A. parasiticus* strains had the most predicted BGCs as NRRL 2999 and NRRL 502 were predicted to contain 80 and 81 BGCs, respectively. Both *A. arachidicola* strains encode more predicted BGCs (76 for IC26645 and 73 for IC26646) than either *A. flavus* strain, but fewer than either *A. parasiticus* strain. Our two *A. flavus* strains, NRRL 1957 and NRRL 502, encode 70 and 71 BGCs, respectively (Fig. 3A). Of these BGCs, 44 (NRRL 1957) and 45 (NRRL 502) were not linked to any known metabolites.

**Figure 3.**
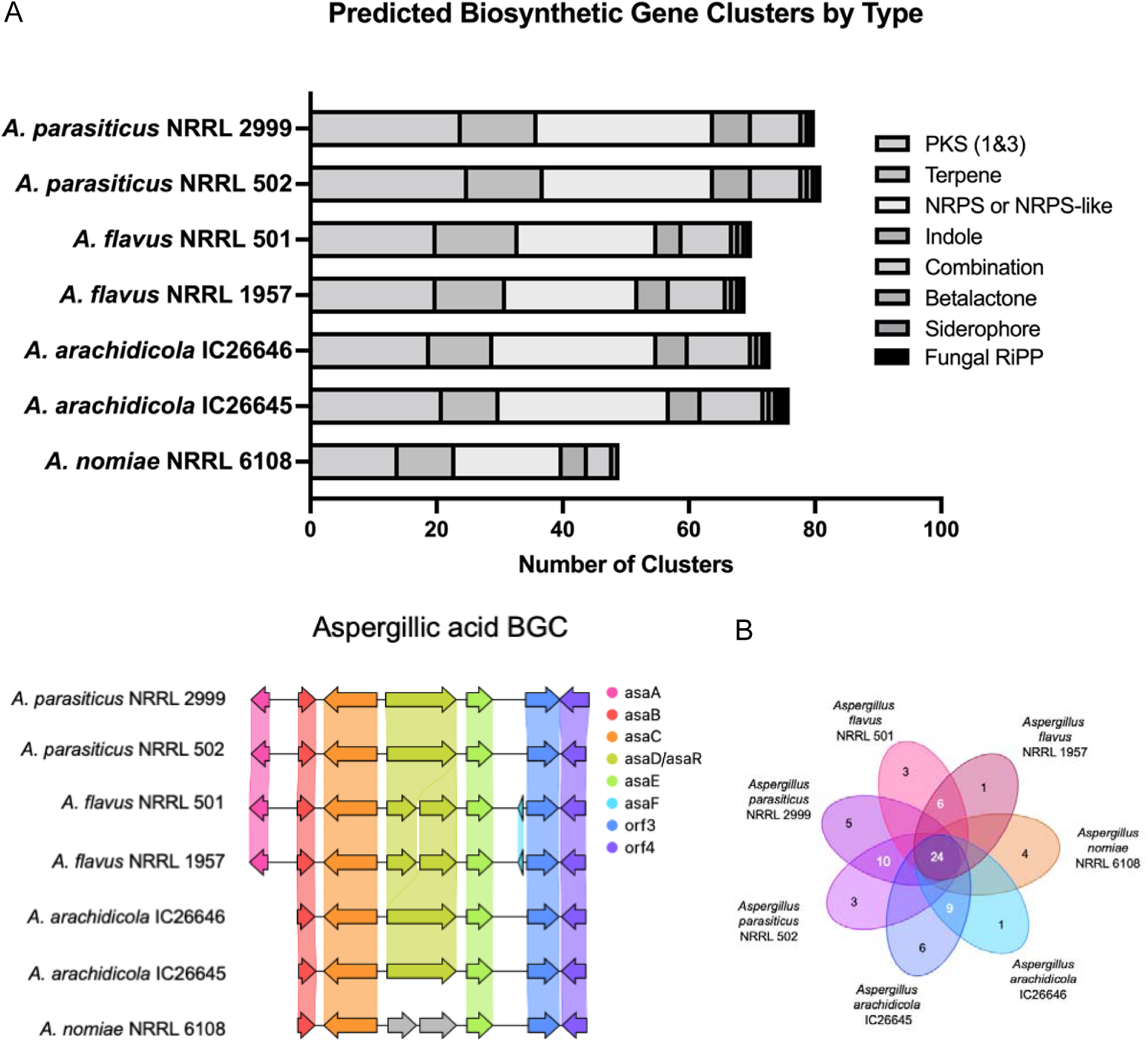
Strains of the same species do not differ substantially in their predicted biosynthetic gene clusters (BGCs). A) Stacked bar plot of predicted BGCs. Each bar adds up to the total number of predicted BGCs, with the type of BGC indicated by color. B) Synteny plot comparison of the aspergillic acid BGC from all seven strains. Arrows represent genes, vertically shaded areas between arrows indicate sequence similarity. C) Diagram of unique (singletons) and shared BGC families, calculated by BiGSCAPE.

**Figure 4.**
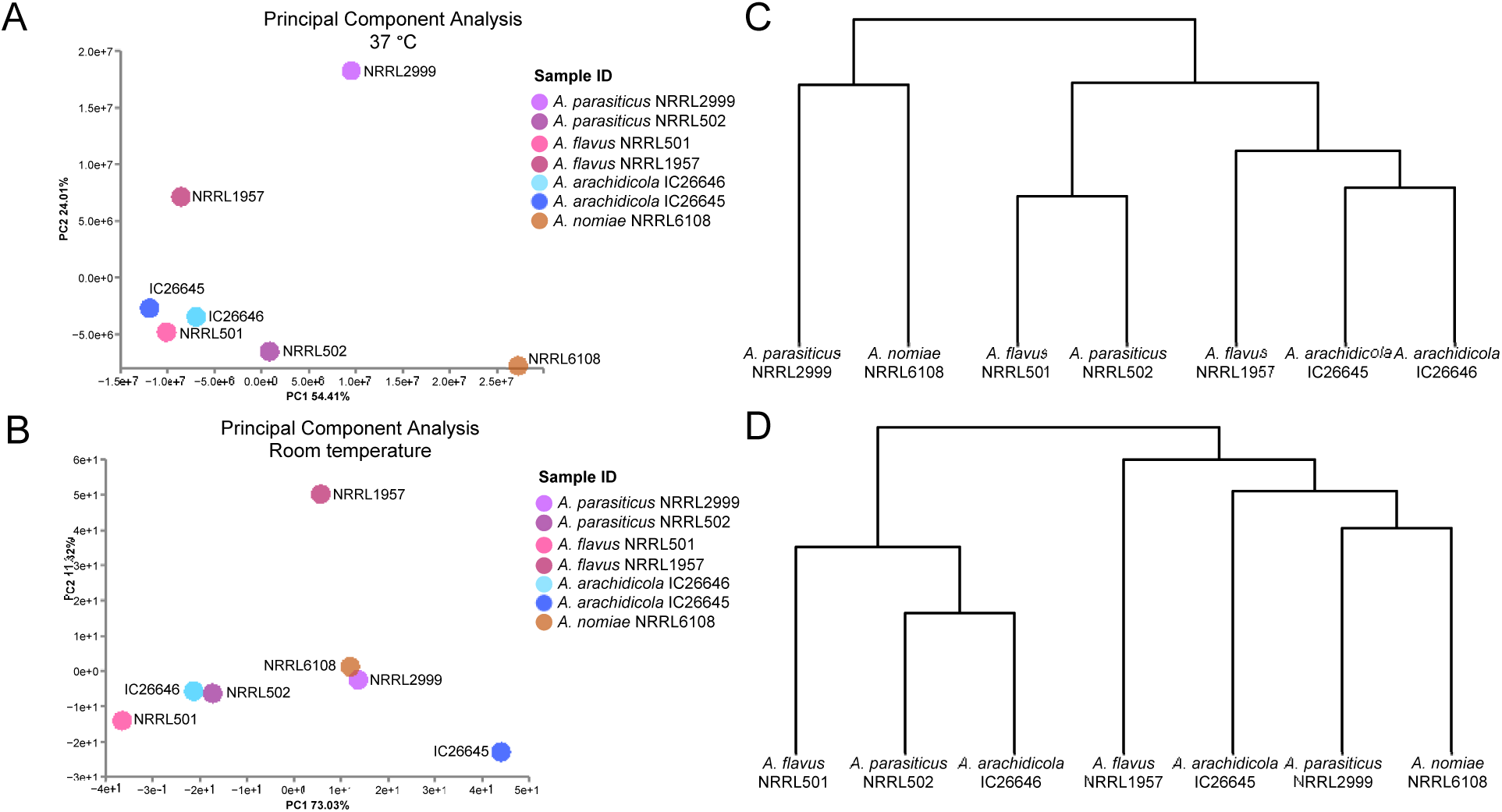
*Aspergillus flavus* strains are more similar at 37°C than room temperature. The metabolomic profiles of *A. flavus* NRRL 501 and NRRL 1957 are almost identical at 37°C showing very similar metabolites in the UPLC-MS analysis; most of the metabolites identified are fatty acids and ergosterol derivatives. In contrast, they are significantly different at room temperature. A) Principal component analysis for all strains at 37°C. Circles represent both presence of metabolites and relative abundance. B) Principal component analysis for all strains at room temperature. Circles represent both presence of metabolites and relative abundance. C) Hierarchical clustering of strains based on metabolite profiles at 37°C. D) Hierarchical clustering of strains based on metabolite profiles at room temperature.

A total of 118 BGC families are present in the seven strains, with 24 families shared by all strains and 7 families unique to the *A. flavus* strains (Fig. 3C). Eight additional BGC families were identified in all strains but *A. nomiae* (Table S1). Our *A. flavus* strains share 58 BGC families (Table S1).

Evaluating the secondary metabolite profiles of the seven fungi at both room temperature and 37°C, and with or without the inclusion of saline, led to several key observations. First, the inclusion or absence of saline had little to no effect on the secondary metabolite profiles, as major differences were not observed between strains grown under the same conditions (i.e., room temperature or 37°C) both with and without physiologic saline (Figures S1 and S2). As such, our analysis of secondary metabolites focused on the impact of temperature. In general, the cultures grown at room temperature contained a high proportion of mycotoxins (i.e., compounds **1**-**32**) or ergosterol derivatives (**33**), whereas the cultures grown at 37°C had a much higher concentration of metabolites derived from fatty acids (i.e., compounds **34**-**68**). Interestingly, the two strains of *A. flavus* (i.e., NRRL 501 and NRRL 1957) did not share any compounds in their secondary metabolite profiles when grown at room temperature. Moreover, other than leporin B and C (**10** and **11**, respectively) and cyclopiazonic acid (**18**), the secondary metabolite profiles of the two strains of *A. flavus* did not overlap with the secondary metabolite profiles of any of the other section *Flavi* species when grown at room temperature. *A. flavus* strain NRRL 501 largely biosynthesized aspergillic acid (**29**) and a variety of related analogues, confirming previous reports (62), whereas *A. flavus* strain NRRL 1957 biosynthesized cyclopiazonic acid (**18**) and range of analogues, such as speradines A (**19**), C (**21**), F (**22**), I (**23**), and H (**24**), and asperorydine H (**25**). Additionally, this strain biosynthesized indol-terpenoids, such as paspaline (**26**) and aflatrem (**27**).

*A. parasiticus* NRRL 2999 and *A. arachidicola* IC26645 had very similar secondary metabolite profiles and are prolific producers of aflatoxins B1 (**1**), B2 (**2**), G1 (**3**) and G2 (**4**), astellolids A (**6**), B (**7**) and D (**8**), and leporins B (**10**) and C (**11**), along with some minor metabolites, such as aspergillicin A (**12**) and phomaligol A (**13**). In contrast, *A. parasiticus* NRRL 502 and *A. arachidicola* IC26646 did not biosynthesize aflatoxins or leporins. However, *A. parasiticus* NRRL 502 produces astellollids A (**6**) and B (**7**) and polyketides like phomaligol A (**13**), phomagilin A (**14**) and aspersitin (**15**), whereas the latter (i.e., *A. arachidicola* IC26646) only produces the 14-deacetyl derivative of astellolide A (**9**) and traces of aspergillicin A (**12**) and phomaligol A (**13**). Furthermore, we observed that *A. nomiae* NRRL6108 biosynthesized a smaller suite of mycotoxins, which is consistent with the smaller number of predicted BGCs in its genome; however, it uniquely produced *O*-methylsterigmatocystin (**5**) and parasiticol (**16**), two metabolites closely related to aflatoxins and likely derived from similar BGCs.

The isolated metabolites at 37°C were largely composed of fatty acids (mostly C_18_ and C_16_ derivatives), particularly the sporogenic PSI (precocious sexual inducer) factors A, B and C (i.e., **41**-**45**). These compounds were found in all the strains grown at 37°C. A few of the mycotoxins were also observed at elevated temperatures, specifically astellolids A (**6**) and B (**7**) in *A. parasiticus* NRRL 2999 and *A. arachidicola* IC26645, and *O*- methylsterigmatocystin (**5**) in *A. nomiae* NRRL 6108. In total, the results showed a surprising amount of heterogeneity of the secondary metabolomic profiles both between species and even between strains of the same species. Structures for all identified metabolites are available in the supplement (Table S2).

### A. flavus strains responded differently to cell wall stress and antifungal drug resistance, but similarly to hypoxia and iron starvation

Hypoxia impacted growth of all strains negatively, but *A. parasiticus* NRRL 2999 was significantly less impacted compared to all other strains (p = 0.0346). Growth of *A. arachidicola* IC26646 was most impacted by the hypoxic environment, although it did not significantly differ from *A. arachidicola* IC26645 or *A. flavus* NRRL 1957. Responses of *A. parasiticus* strains to hypoxia were significantly different (p = 0.027), whereas observed differences between strains of *A. flavus* and between *A. arachidicola* strains were not (Fig. 5A).

**Figure 5.**
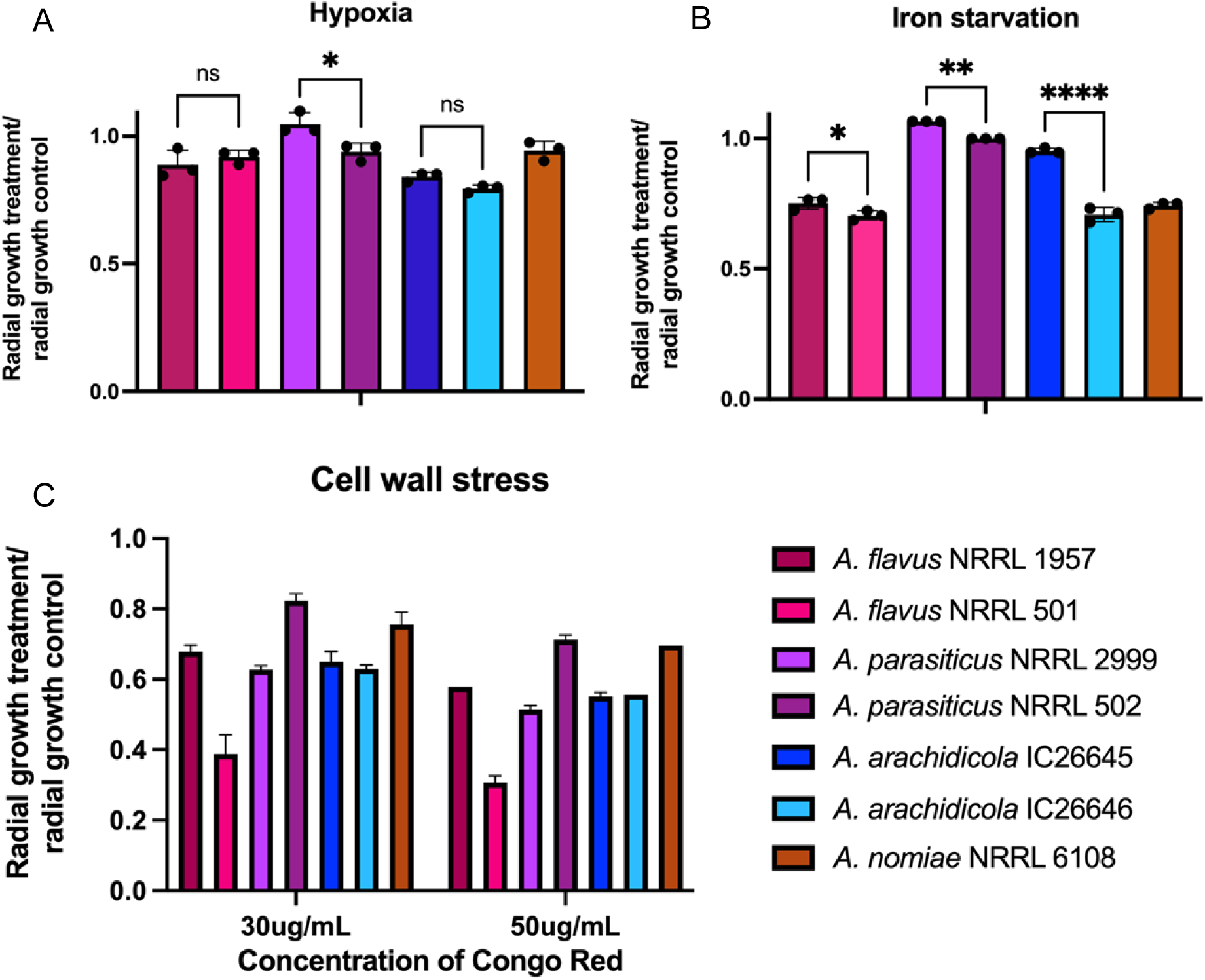
Iron starvation and cell wall stress impact growth of *A. flavus* strains differently, but hypoxic conditions impact *A. flavus* strains similarly. For each of the seven *Aspergillus* strains, radial growth is expressed as a ratio, dividing colony radial diameter (cm) of growth in the stress condition by colony radial diameter in the control (solid minimal media) condition. Not all significant comparisons are shown. A) Hypoxic stress was induced by incubating plates in 1% O_2_ and 5% CO_2_. Statistical significance of growth differences among species is primarily driven by growth of *A. parasiticus* NRRL 2999, which was significantly less impacted by hypoxia than other strains. B) Iron starvation was induced through growth on iron-depleted substrate in the presence of gallium. All species with multiple strains exhibited strain heterogeneity, and *A. parasiticus* grew significantly better in iron starvation conditions than other species. Other species comparisons between species were nonsignificant. C) Cell wall perturbation was induced by adding Congo red to the medium. *A. flavus* NRRL 501 was most impacted by Congo red at both concentrations, and *A. parasiticus* NRRL 502 the least. At both concentrations, strains of *A. flavus* had significantly different responses to cell wall stress (p < 0.0001). Cell wall stress also impacted *A. parasiticus* strains differently (p < 0.0001). Strains of *A. arachidicola* did not have significant growth differences. Statistics: ANOVA, ns = not significant, * p ≤ 0.05, ** p < 0.005, *** p < 0.0005

Under iron starvation conditions (growth in iron-depleted media), *A. parasiticus* NRRL 2999 was least impacted and grew slightly better than in iron-supplemented media. Growth of *A. flavus* NRRL 501 was most impacted by the lack of iron (Fig. 5B). *A. flavus* strains differed significantly from *A. parasiticus* strains under iron starvation conditions (p < 0.0001, one-way ANOVA) and from one another (p = 0.0398). Growth differences of the two *A. parasiticus* strains under iron starvation conditions were also statistically significant (p = 0.0030), as were dissimilarities in growth rates of the two *A. arachidicola* strains (p < 0.0001). *A. flavus* NRRL 1957 was more sensitive to oxidative stress compared to other strains (p = 0.0003).

Strains of *A. flavus* had varying responses to cell wall stress, with a stronger impact of Congo red on the growth of NRRL 501 than NRRL 1957 (p < 0.0001, two-way ANOVA). *A. flavus* NRRL 501 was significantly more sensitive to cell wall stress than any other strain (p < 0.0001). *A. parasiticus* strains also exhibited different growth rates in the presence of Congo red, with NRRL 2999 growing less than NRRL 502 (p < 0.0001). Growth of the two *A. arachidicola* strains was not significantly different from each other or from *A. flavus* NRRL 1957 for either concentration of Congo red (Fig. 5C).

*Aspergillus parasiticus* NRRL2999 had the lowest minimum inhibitory concentration (MIC) for amphotericin B (0.5 µg/ml), with all other strains requiring a higher dose to inhibit growth (1 µg/ml). Both *A. arachidicola* strains and *A. nomiae* NRRL 6108 had a higher MIC for voriconazole, with *A. parasiticus* and *A. flavus* strains being more susceptible (Table 5).

**Table 5.**
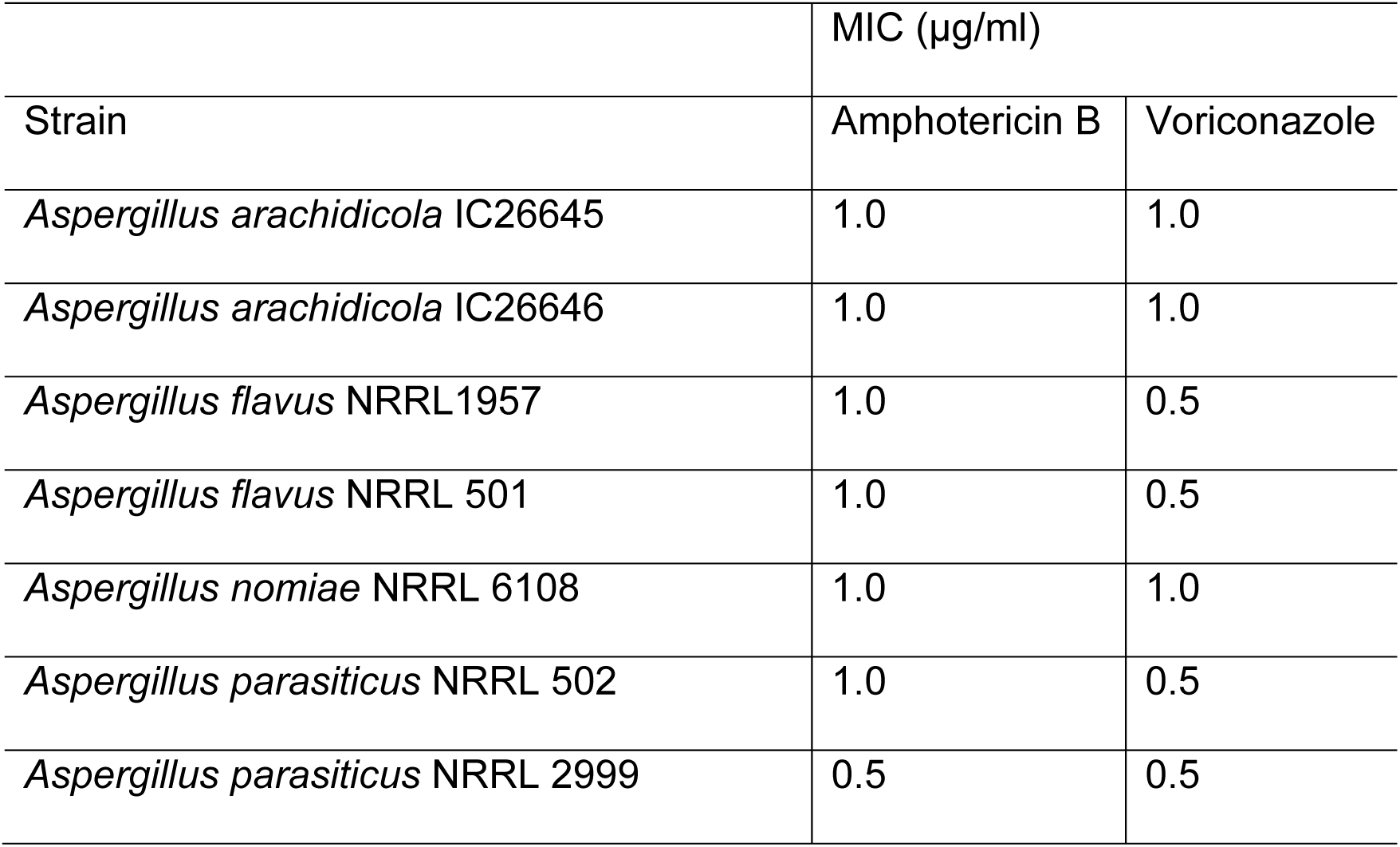
Minimum inhibitory concentrations of the antifungal drugs amphotericin B and voriconazole for seven strains of four *Aspergillus* species

| Strain | MIC (µg/ml) |  |
| --- | --- | --- |
|  | Amphotericin B | Voriconazole |
| *Aspergillus arachidicola* IC26645 | 1.0 | 1.0 |
| *Aspergillus arachidicola* IC26646 | 1.0 | 1.0 |
| *Aspergillus flavus* NRRL1957 | 1.0 | 0.5 |
| *Aspergillus flavus* NRRL 501 | 1.0 | 0.5 |
| *Aspergillus nomiae* NRRL 6108 | 1.0 | 1.0 |
| *Aspergillus parasiticus* NRRL 502 | 1.0 | 0.5 |
| *Aspergillus parasiticus* NRRL 2999 | 0.5 | 0.5 |

### A. flavus is not significantly more virulent than related, non-pathogenic species in an invertebrate model of fungal disease

Using an invertebrate model of fungal disease, we evaluated virulence for all strains. *A. fumigatus* virulence assays typically use concentrations of 1 x 10^6^ conidia (asexual spores) to inoculate *G. mellonella* larvae (64, 65). When we inoculated larvae with section *Flavi* species at 1 x 10^6^, all animals died within 2 days. Previous studies provide evidence that *A. flavus* kills *G. mellonella* larvae faster than *A. fumigatus* and therefore requires a lower inoculum (63). At a lower concentration inoculum, 1 x 10^4^ conidia (spores), the larvae survived longer and differences among strains were apparent.

We found that strains of the same species varied widely in their virulence profiles, and that strains of the pathogen *A. flavus* were not more virulent than strains of the non-pathogenic species (Fig. 6). Using the 1 x 10^4^ concentration of spores, *A. parasiticus* NRRL 2999 killed the fewest animals and was not statistically different from the PBS control injections. In contrast, *A. arachidicola* IC26646, *A. flavus* NRRL 1957, and *A. parasiticus* 501, killed all inoculated moth larvae by day three. Of note, the three most virulent strains were all from different species, as were the three least virulent strains; strains of the same species killed larvae at significantly different rates (Fig. 6).

**Figure 6.**
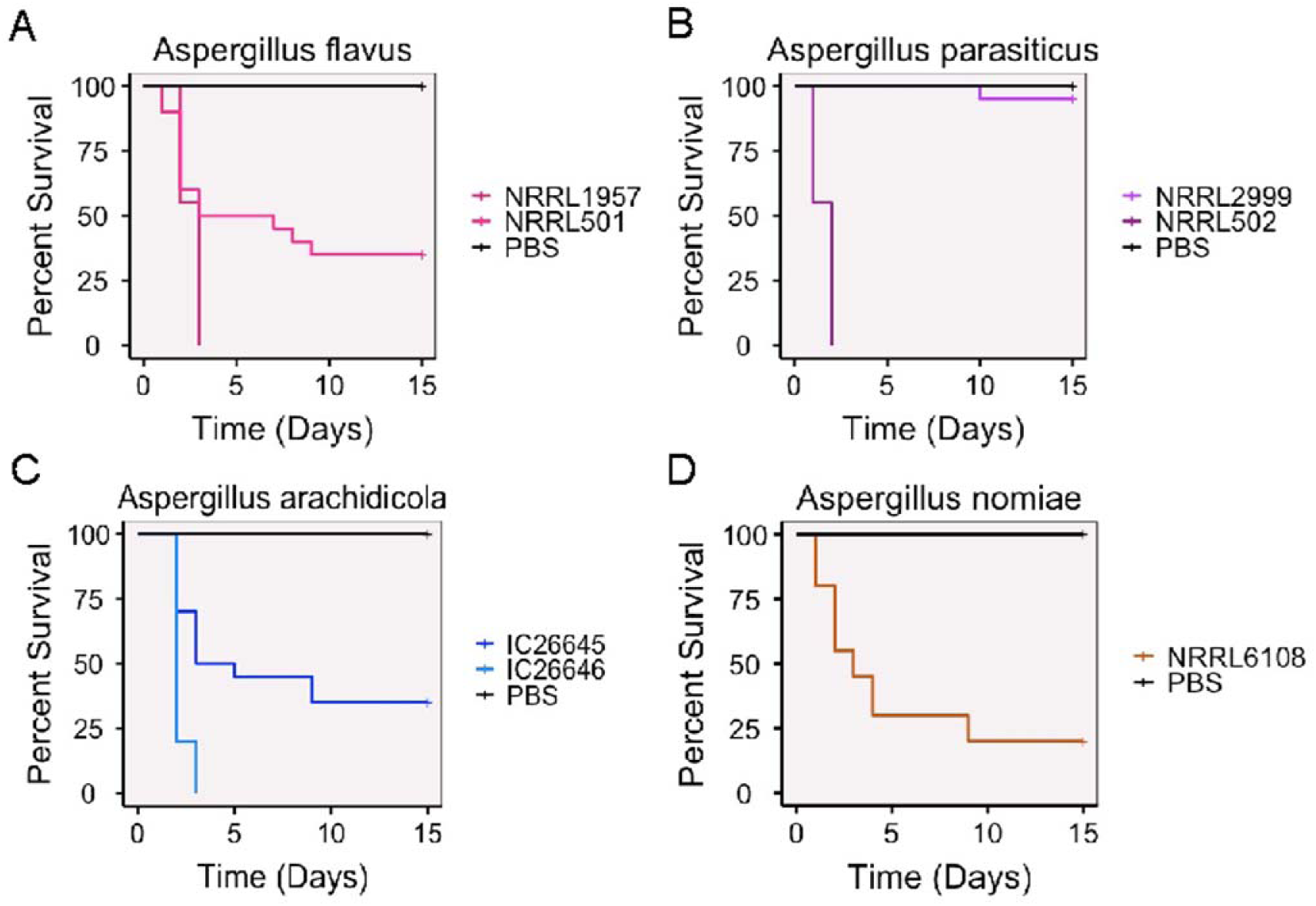
*Aspergillus flavus* is not significantly more virulent than related non-pathogenic species in an invertebrate model of fungal disease. Cumulative survival of *Galleria mellonella* larvae inoculated with 1 x 10^4^ asexual spores (conidia) of an *Aspergillus* strain or a PBS control. A) Survival for larvae inoculated with either *A. flavus* NRRL 1957 or NRRL 501. All pairwise comparisons between the two strains and the control group were statistically significant. B) Survival for larvae inoculated with either *A. parasiticus* NRRL 2999 or NRRL 502. *A. parasiticus* NRRL 2999 was not statistically different from the control group, but NRRL 502 was statistically different from both the control and NRRL 2999. C) Survival for larvae inoculated with either *A. arachidicola* IC26645 or IC26646. All pairwise comparisons between the two strains and the control group were statistically significant. D) Survival for larvae inoculated with *A. nomiae* NRRL 6108. NRRL 6108 was statistically different from the control.

Flavi *strains infect eyes as well as* A. fumigatus *in a murine model of fungal keratitis* Finally, we used a mouse model of keratitis to compare one strain from each of the four different *Aspergillus* species at 24, 48, and 72 hrs post infection (Fig. 1A). We included *A. fumigatus* Af293 along with the three *Flavi* species (*A. arachidicola* IC26646, *A. flavus* NRRL 1957, and *A. parasiticus* NRRL 2999) to benchmark the virulence of *Flavi* species against the reference strain for section *Fumigati*. We observed significantly lower disease severity (Fig. 7B) and corneal thickness (Fig. 7C) in *A. arachidicola* IC26646 infections compared to the other species at 48 hrs post infection, but this difference was not significant at 72 hrs. Fungal burden was not significantly different between the four species (Fig. 7D).

**Figure 7.**
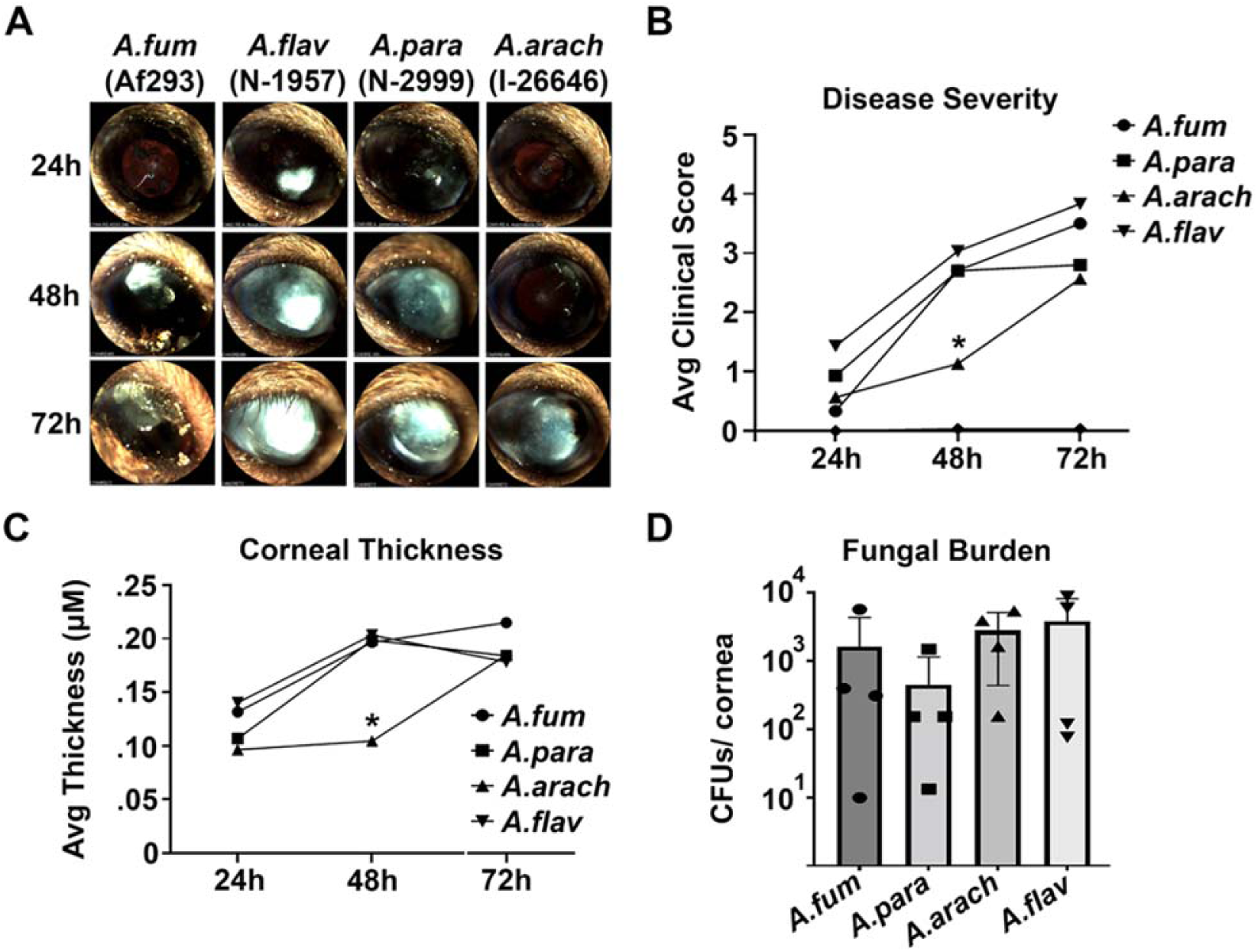
*Flavi* species infect eyes as well as *A. fumigatus* in a murine model. A) Slit-lamp images of a representative animal for each infection group. B) Clinical score analysis of all slit-lamp images reveals reduced disease severity in the *A. arachidicola* group at 48h post-infection (n=5/group). C) Corneal thickness measured by optical coherence tomography similarly reveals reduced structural alteration in the *A. arachidicola* group at 48h (n=5). D) Colony forming unit (CFU) analysis on resected corneas reveal indistinguishable fungal burden at 72h post-infection between infection groups. *A.arach* = *A. arachidicola* IC26646, *A.flav* = *A. flavus* NRRL 1957, *A.fum* = *A. fumigatus* Af293, *A.para* = *A. parasiticus* NRRL 2999.

## DISCUSSION

With the goal of studying the evolution of pathogenicity in section *Flavi*, we examined the genomes, chemotypes, and phenotypes of four closely related species: the major pathogen *A. flavus* and three related non-pathogenic species *A. arachidicola*, *A. parasiticus*, and *A. nomiae.* We observed similarities and differences between *A. flavus* and three non-pathogenic relatives, including shared gene content, variable production of secondary metabolites, and virulence in two animal disease models.

Genomic content was highly similar between strains of the same species, with few strain-specific protein families identified. Predicted BGCs were similar between strains of the same species, with 21 shared by all strains. Georgianna et al. (64) previously predicted 55 biosynthetic gene clusters (BGCs) from *A. flavus* NRRL 3357. Of these, 14 BGCs have been linked to a specific metabolite. Additional clusters producing kojic acid, aflavinines, apseripin-2a, and ustiloxins were later identified. Recently, Drott et al. (65) studied 94 different *A. flavus* strains and identified 92 unique BGCs. Our *A. flavus* strains were predicted to contain 70-71 BGCs, well within the expected range, although over half of the predicted BGCs in each strain have not been linked to known metabolites.

Among the species with multiple strains included, the two *A. parasiticus* strains shared the fewest species-specific protein families (1, 230), whereas pairs of strains from *A. arachidicola* and *A. flavus* shared over 3,000 species-specific protein families between strains, around 20% of the total number of protein families for each species. Our results within *Flavi* contrast with observations within *Fumigati*, which show a lower proportion of species-specific genes for *A. fumigatus* compared to related, non-pathogenic species (66). Specifically, when strains of *A. fumigatus* were compared to other *Fumigati* species, only 72 families were identified as unique to *A. fumigatus* (66); in contrast, we observed over 3,000 unique families in *A. flavus*. Although section *Flavi* species are known to have larger genomes and encode more predicted proteins than *A. fumigatus* (29), this does not explain the huge difference in number of species-specific protein families between *A. fumigatus* and *A. flavus*, and further emphasizes that the two sections are quite distinct in their genomic compositions.

Resistance to antifungal drugs was also similar between strains of the same species and within previously reported ranges for *A. flavus* (67). Our *A. flavus* and *A. parasiticus* strains were more susceptible to amphotericin B than clinical strains of the same species from Brazil (68), and strains from all four species had similar susceptibility to voriconazole as clinical strains from Brazil (69, 70). As expected, our *Flavi* strains were generally more resistant to amphotericin B than to voriconazole.

Several traits examined revealed heterogeneity between strains of the same species, indicating diversity within each species as well as among species. Our two *A. flavus* strains, NRRL 501 and NRRL 1957, for example, did not produce any of the same secondary metabolites when grown at room temperature, and only a handful of compounds produced by the two strains at room temperatures overlapped with those produced by strains of other species. We found that our *A. parasiticus* strains produced aflatoxin at higher levels than other species, including noted aflatoxin-producer *A. parasiticus* NRRL 2999, although one strain of *A. flavus*, NRRL 1957, had previously been characterized as negative for aflatoxin (70). At the temperature of the human body, which better models infection-relevant conditions, several compounds were produced by all seven strains. These include fatty acids such as palmitic acid, which has been implicated in inflammation (71), and precocious sexual inducer factors. Interestingly, our two *A. flavus* strains had highly similar chemical profiles at 37°C.

Strains of the same species also responded differently to environmental stressors in growth assays, including cell wall stress and iron starvation, which may affect a strain’s fitness within a human host. For example, the ability to tolerate cell wall stress (induced by Congo red) correlated with higher virulence in an invertebrate model, as observed in *A. flavus* NRRL 1957 and *A. parasiticus* NRRL 502.

Additionally, in *Galleria mellonella*, neither *A. flavus* strain was the most virulent, and all species were able to infect and kill at least some larvae, although *A. parasiticus* NRRL 2999 killed only 10% of larvae. As observed previously, the concentration of asexual spores required to kill all larvae within 24 hrs is orders of magnitude lower for infection with *Flavi* species compared to *Fumigati* species (63), suggesting higher virulence in the *Flavi* species. In the most virulent *Flavi* strains (*A. parasiticus* NRRL502, *A. flavus* NRRL1957, and *A. arachidicola* IC26646), death of all larvae occurred within 72 hrs. A previous study using a lower concentration inoculum (1 x 10^3^ spores) saw 100% mortality of larvae in the first 48 hrs for *Galleria* infected with *A. flavus* (72). However, in another study of *A. flavus* strains, also with a concentration of 1 x 10^3^ spores, 10% or more of the larvae were alive after 72 hrs for all strains (73). Few studies of non-*A. flavus* section *Flavi* species include virulence assays in *Galleria*, presumably due to their status as non-pathogenic, preventing direct comparison between our results and previous studies for species other than *A. flavus*. As a known entomopathogen (74), virulence by *A. flavus* in an invertebrate model may be correlated with insecticidal traits rather than (or in addition to) human pathogenicity.

To examine virulence in a mammalian model of fungal keratitis, we infected eyes of mice with one strain each of *A. flavus*, *A. arachidicola*, and *A. parasiticus*, along with one *A. fumigatus* strain. We observed faster colonization of mice by *A. flavus* than *A. fumigatus* at 24 hrs, although fungal burden and disease severity was consistent for all species by 72 hrs. *A. arachidicola* IC26646 did not infect as quickly, with significantly lower disease severity observed at 48 hrs post infection, but caught up by 72 hrs post infection. Mouse models of disease are a useful tool for understanding disease progression and characterizing differences among strains, but previous studies have induced systemic infections leading to mouse death (75). Systemic infection studies have shown *A. flavus* to be more virulent than *A. fumigatus* (22, 75), despite higher rates of aspergillosis caused by *A. fumigatus*. Interestingly, we observed no difference in disease severity at 72 hrs between the two species in our keratitis model. The similarity of secondary metabolites profiles for strains at 37°C, coupled with the ability of all species to establish infections in both the *Galleria* and murine models, leads us to believe that infection may be more highly dependent on the characteristics of the individual strain rather than species, as observed in other pathogens from *Aspergillus* (76). We intend to compare additional strains of both *A. flavus* and *A. fumigatus*, including patient-derived clinical strains, to examine differences in virulence and disease progression of *A. flavus* and *A. fumigatus* in our keratitis model. It remains an open question whether patient-derived clinical strains of *A. flavus* differ from the environmental strains available from culture collections, such as the two strains included in this study, and we hope to explore this dimension of pathogenicity in future studies.

In summary, by examining genomic, chemical, and phenotypic variation within and between closely related pathogenic and non-pathogenic *Aspergillus* section *Flavi* species, we show that species considered non-pathogenic infect at the same rate as the pathogen *A. flavus* in both invertebrate and murine models of disease, and that strain-level differences may play a major role in infection.

## Data Availability Statement

Sequencing data and genome assemblies associated with this project is listed under BioProject <u>PRJNA824811</u>. Raw reads are available through the NCBI sequence read archive under accessions <u>SRR19347655</u> (*Aspergillus arachidicola* IC26646), <u>SRR19347656</u> (*Aspergillus arachidicola* IC26645), <u>SRR19347505</u> (*Aspergillus flavus* NRRL 1957), <u>SRR18725159</u> (*Aspergillus flavus* NRRL 501), <u>SRR19347653</u> (*Aspergillus parasiticus* NRRL 2999), <u>SRR19347654</u> (*Aspergillus parasiticus* NRRL 502), and <u>SRR19369914</u> (*Aspergillus nomiae* NRRL 6108).

1H and ^13^C NRM data for compounds **1**-**68** will be submitted to the Natural Products Magnetic Resonance Database (https://np-mrd.org/) upon manuscript acceptance. Supplemental tables and figures are available on FigShare (https://figshare.com/s/fa424e3b0be171d5d7ba).

## Acknowledgements

We thank the Rokas lab, and in particular Dr. Matthew Mead, for helpful discussion and feedback. Research in A.R.’s lab is supported by grants from the National Science Foundation (DEB-2110404), the National Institutes of Health/National Institute of Allergy and Infectious Diseases (R56 AI146096 and R01 AI153356), and the Burroughs Wellcome Fund.

